# Beyond the circuit architecture : attractor dynamics reveals the mechanism of improved performance in decision-making and working memory

**DOI:** 10.1101/2022.06.11.495775

**Authors:** Han Yan, Jin Wang

**Affiliations:** State Key Laboratory of Electroanalytical Chemistry, Changchun Institute of Applied Chemistry, Chinese Academy of Sciences, Changchun, Jilin, 130022, P.R. China; Department of Chemistry and Physics, State University of New York at Stony Brook, Stony Brook, NY 11790, USA

**Keywords:** decision-making and working memory, attractor landscape, circuit architecture, non-equilibrium

## Abstract

Understanding the underlying mechanisms of cognitive functions such as decision-making(DM) and working memory(WM) is always one of the most essential concerns in modern neuroscience.Recent experimental and modelling works suggest that decision-making is supported by the selective subnetwork of inhibitory neurons, rejecting the previously proposed circuit mechanisms assuming a single non-selective pool of inhibitory neurons. The mechanism underlying decision-making and working memory functions based on such circuit architecture is still unclear. Here we applied a general non-equilibrium landscape and flux approach to a biophysically based model that can perform the decision-making and working memory functions. The quantified attractor landscapes reveal that the accuracy in decision-making can be improved due to the stronger resting state in the circuit architecture with selective inhibition, while robustness of working memory against distractors is weakened, which implies a trade-off between DM and WM. We found that the presence of a ramping non-selective input during the delay period of the decision-making tasks can serve as a cost-effective mechanism of temporal gating of distractors. This temporal gating mechanism, combined with the selective-inhibition circuit architecture, can support a dynamical modulation for emphasizing the robustness or the flexibility to incoming stimuli in working memory tasks according to the cognitive task demands. These mechanisms can also achieve an optimal balance in the trade-off between DM and WM. Our approach can provide a global and physical quantification which helps to uncover the underlying mechanisms of various biological functions beyond the circuit architectures.

---

The brain, as a complex dynamical system, can perform various physiological and cognitive functions(1–6). The computational abilities involved in these functions emerge as collective properties of neural circuits in which enormous number of neurons interact with each other via excitation or inhibition. Decision-making(DM) and working memory(WM) are fundamental building blocks of cognition, and great efforts in both theoretical and experimental fields have been made to uncover the underlying mechanisms of the corresponding neural circuits, e.g. how excitatory and inhibitory neurons coordinate during DM and WM.

DM task such as perceptual discrimination or target selection for motor response is associated with ramping activity of neurons in the parietal, prefrontal or premotor cortex, reflecting the accumulation of evidence(3, 7–10). The decision choice made in the DM task or other sensory information is sometimes required to be actively held for a short period of times even after the withdrawal of the external stimulus. Such WM function is associated with the stimulus-selective persistent activity, which is commonly observed in the same (parietal, prefrontal, premotor) cortical areas(3, 8, 10–12). These experimental recordings suggest that the neural circuits endowed with persistent activity may also be capable of performing stimulus integration and categorical decision choice. Moreover, DM and WM may share similar circuit mechanisms. Recent theoretical works show that the characteristic neural activity of DM and WM can be successfully described by the attractor network framework(3, 10, 13, 14). In the attractor networks that are capable of performing both WM and DM functions, there are two or more populations, which have self-excitation and are selective to different stimuli. These excitatory populations interact with each other through a common pool of inhibitory population(3, 10, 13, 14). The neural circuits with such self-excitation and mutual inhibition architecture can generate the stimulus-selective persistent activity pattern for WM(11, 12, 15) and also ramping, categorical and winner-take-all dynamics for perceptual DM(10, 16).

Inhibitory neurons, which serve as an essential component in these models, are usually provided by a single pool of non-selective neurons lacking connection specificity(3, 10, 13, 14). However, recent studies find that both the excitatory and inhibitory neurons in decision circuits are selective, which suggest that specific functional subnetworks exist within inhibitory populations, just as the situation in excitatory populations(5, 17). These results raise the question that among possible circuit architectures that can support DM, why such particular circuit architecture with functionally specific connected subnetworks(selective inhibition) is formed during learning? Could such selective-inhibition architecture improve the performance of DM and further the corresponding WM? If the selective-inhibition architecture couldn’t protect memory from interference during DM, is there any additional mechanism beyond the circuit architecture?

To address these questions, we tried to investigate the mechanisms underlying the DM and WM functions based on the attractor network framework. The attractor network models can reproduce most of the psychophysical and physiological results in the DM and WM tasks(8, 18). Mostly, attractor dynamics provide a natural candidate mechanism for mnemonic persistent neural activity and winner take-all competition leading to a categorical choice. Noise is ubiquitous in the brain, e.g. intrinsic random fluctuations caused by highly irregular spiking activity within cortical circuits and the stochastic external inputs. In such noisy systems, stochastic transitions between different states(attractors) represent switching between perceptual or physiological states. Quantifying the underlying attractor landscapes can provide a global picture of noisy neural systems, including global stability of functional states and transitions between them. Nevertheless, the attractor landscape described extensively in previous works are usually not given explicitly but only as illustrations(3, 10, 13). Inspired by the thermodynamics and statistical mechanics, physicists introduced the concepts of “internal energy”, “free energy” and “entropy” to describe the collective properties of complex systems, such as neural circuits and proteins(1, 19, 20). Hopfield pioneeringly explored the computational properties of neural circuits, e.g. memory storage and retrieval, by constructing an “energy function”(1, 19). However, the energy function in the original Hopfield model can only be constructed for symmetrical neural circuits, and the original Hopfield model fails to apply for more realistic biological neural circuits under asymmetrical connections. Furthermore, neural circuits, as other biological systems, consume energy to perform different vital functions(21, 22). The essential role of the energy supply in cognitive processes such as DM and WM has yet to be fully clarified.

Here we applied a general non-equilibrium landscape and flux approach(14, 23, 24) to a biophysically based model that can perform DM and WM functions(3, 10). We probed the mechanisms of these cognitive functions through quantifying the underlying attractor landscapes which determine the behavioral responses of the system in DM and WM tasks. We found that the circuit architecture with selective inhibition results in stronger resting state, which improves the accuracy of DM. Nevertheless, the weaker decision states in this circuit structure lead to less stable working memories against distracting stimuli. Our results show that presenting a ramping non-selective input in the early-delay period can serve as a temporal modulation mechanism that enhances the robustness of WM with less energy consumption. Moreover, our results predict that this temporal gating mechanism combined with the selective-inhibition architecture may support a dynamical modification of emphasizing the robustness or flexibility to incoming stimuli in WM tasks according to the cognitive task requirement. Our approach provides a new paradigm for exploring the underlying mechanisms of various cognitive functions.

## Results

### Influence of the circuit architecture on the DM and WM functions

Both the posterior parietal cortex (PPC) and prefrontal cortex (PFC) are key nodes that can exhibit characteristic neural activity of cognitive functions such as DM and WM(25–27). The perceptual decision-making behavior and working memory function in these cortical areas were well explored through the classic random-dot motion (RDM) discrimination tasks(28–31). In such experimental paradigm, monkeys are trained to judge the direction of motion of random dots, and their choice responses are indicated by a rapid saccadic eye movement. In a delayed response version of the task(8), the monkey is required to withhold the response across a delay period, which means its choice must be maintained actively in working memory. To account for these functions, previous modeling works show that DM-related ramping dynamics and winner-take-all competition while WM-related persistent activity can be performed in neural circuits with the architecture that pools of stimulus-selective excitatory neurons compete with each other through feedback inhibition from a common pool of non-selective inhibitory neurons(Fig.1a)(3, 10, 13, 14, 16). However, Recent experimental results argue against circuit architectures assuming non-selective inhibitory neurons and suggest that selective subnetworks, as the ones in the excitatory population, emerge simultaneously within the inhibitory population during learning(Fig.1b)(5, 17). To study why such circuit architecture is chosen from many different alternatives, we explored a biologically realistic attractor model to test how different circuit architectures influence the performance in DM and further WM tasks.

**Fig.1.**
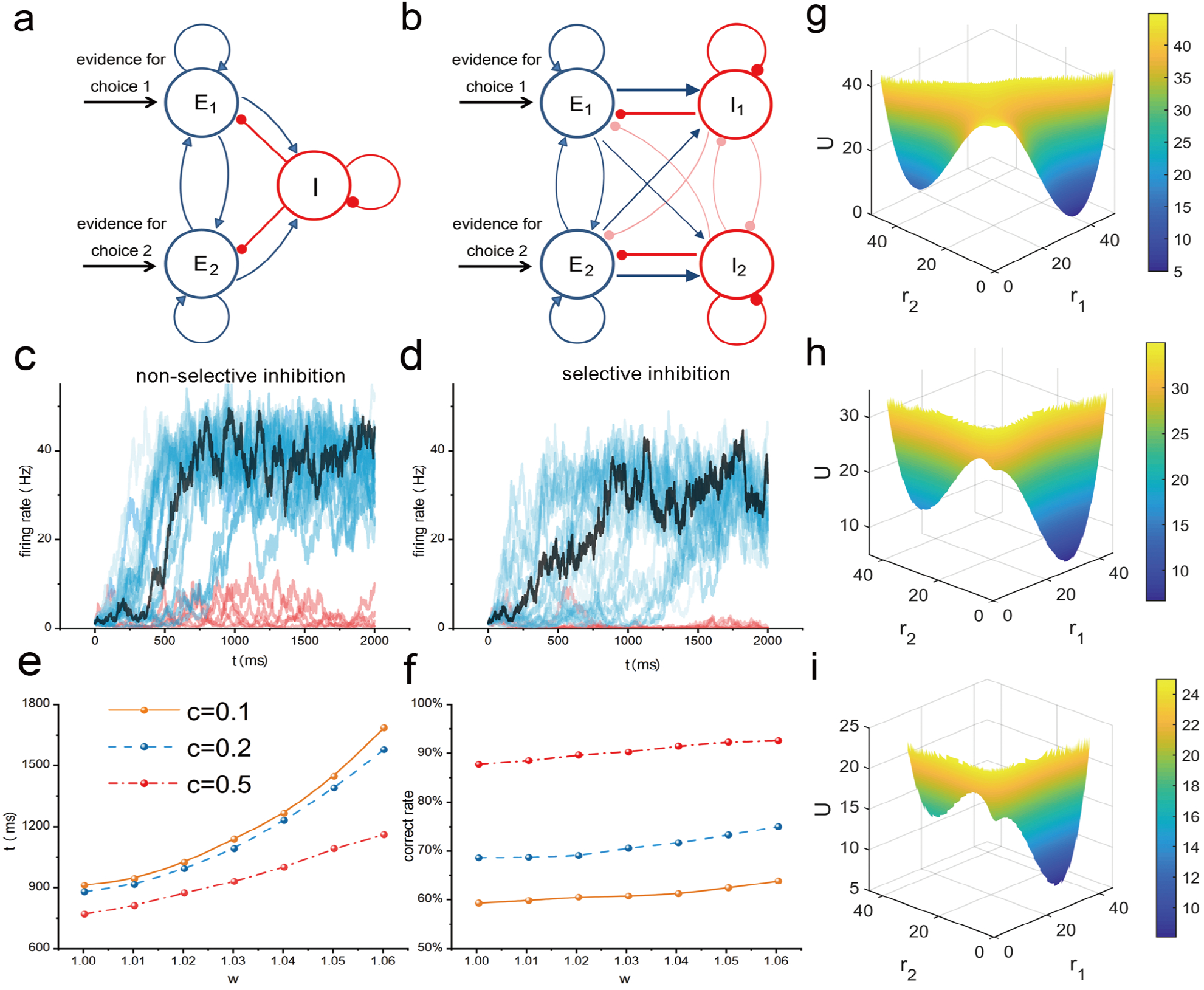
Selective subnetwork of inhibitory neurons enhances the accuracy in decision-making. (a) The diagram of the circuit model with non-selective inhibitory neurons. (b) The circuit architecture with selective subnetwork of inhibitory neurons. The control parameter *w* can indicate the degree of selectivity of the inhibitory neurons. Larger *w* implies the inhibitory neurons more more selective. (c-d)Neural activities during DM for different circuit architectures with non-selective and selective inhibitory neurons, where *w* = 1, 1.06, respectively. The blue curves represent the activities of the excitatory neural population 1 for correct trails, while red curves for the error trails. The black trajectories mark the firing rate of population 1 for the trial with median decision time. (e-f) The average decision times and correct rates in DM with varied architectures that indicated by the parameter *w*. For the generality of the results, we explored the DM tasks with different difficulties(stimulus contrast *c* = 0.1, 0.2, 0.5, respectively). Each data point in Figs.3(e-f) are averaged over 30000 trials. The circuit architecture with selective subnetwork of inhibitory neurons(larger *w*) results in longer decision time and higher accuracy in DM. The underlying mechanism can be reflected from the attractor landscapes showing that the selective-inhibition architecture leads to a stronger resting state, which extends the time of integration of evidences((g-i) with *w* = 1.01, 1.04, 1.06,respectively and the nonzero contrast *c* = 0.1).

In the attractor network framework, a decision is made when the system, in the presence of an external stimulus, is driven from the resting state to one of the competing attractor states corresponding to different choices(10, 13). These choice attractors associated with the stimulus-selective persistent activities can be maintained even in the absence of the stimulus. Hence, the network can subserve both decision-making computation and working memory. Here we use a reduced version of the spiking neuronal network models comprised of integrate-and-fire types through a mean-field approach (3, 10, 32), which can reproduce most of the psychophysical and physiological results in delayed response DM tasks(8, 10, 18). In this model, the two excitatory neural populations (selective for rightward and leftward motion directions) receive the external inputs from visual motion stimulus. These inputs are linear functions of the contrast *c*(see details in the method section, the percentage of random dots moving coherently), which sets the bias of the input for one population over the other depending on whether the motion stimulus is in the preferred or non-preferred (null) direction of the cell. For a zero-contrast stimulus(*c* = 0), the two excitatory populations receive equal input. The choice is difficult to be made, where the random noise plays the dominating role. In a high-contrast condition, there is a larger difference in the external input currents to the two excitatory populations. This implies that it is easier to make a decision. In contrast to the assumption of non-selective inhibition in previous theoretical models(Fig.1a), recent studies suggested that the subnetworks of selective inhibitory neurons support decision-making(17). As shown in Fig.1b, we model this circuit architecture assuming selective inhibitory neurons through dividing the inhibitory neurons into two sub-populations(*I*_1_ and *I*_2_), each connected preferentially to one excitatory pool(*E*_1_ or *E*_2_). We use a control factor *w* to modulate the connection strengths between the excitatory and inhibitory sub-populations. The connection strengths within the population 1 or 2 are larger by the factor *w*, while across-population(between 1 and 2) connection strengths are smaller by a factor of *w*_(details in the materials and methods section). Therefore, larger *w* implies the sub-populations are more selective while *w* = 1 corresponds to the completely non-selective case. In this model, a nonzero contrast stimulus suggests that the excitatory population *E*_1_ receives larger external input. A correct choice is reflected in the ramping neural activity of population *E*_1_ to a high-activity state while the population *E*_2_ to a low-activity state. As shown in Figs.1(c-d), the circuit structure indicated by the factor *w* plays a crucial role in determining the time integration in perceptual decisions, here with a contrast stimulus *c* = 0.2. With the non-selective structure(*w* = 1), the neural activity of the population *E*_1_ ramps relatively faster. With a selective structure(*w* = 1.06), the timescale of integration is relatively slower. Easier tasks indicated by the larger contrast *c* always correspond to shorter decision time and higher accuracy. However, for a specific contrast condition, there is a tradeoff between the decision time and the accuracy, which is known as the speed-accuracy tradeoff(SAT) in DM(33, 34). Fig.1(e) depicts the average decision time increases as the factor *w* is increased in different contrast conditions. The longer integration times in decision processes are accompanied by higher accuracy(Fig.1(f)).

To investigate the underlying mechanisms for enhanced accuracy that results from the signal-selective architecture, we quantified the attractor landscapes that characterize the DM function with varied *w*. As shown in Fig1.(g), there are two biased basins of attraction(decision attractors) corresponding to the two choices in the presence of the nonzero-contrast stimulus. The attractor corresponding to the correct choice is stronger, while the other one is weaker. The system initiates at the resting state between the two decision attractors before the onset of the external stimuli. Once the external stimulus is presented, the resting state loses its stability and the system will go “down hill” to one of the two decision attractors. As the control parameter *w* is increased, the inhibitory neurons become more selective. This results in a stronger resting state with local minimum of the potential landscape in between the two competing decision attractors(Figs.1(g-i) with *w* = 1.01, 1.04, 1.06, respectively). The system has to go across a barrier to reach a decision state, therefore longer decision time is required. Meanwhile, the slower integration of evidence reduces the errors that result from the random noise. Our results raise the possibility that the brain chooses the higher-accuracy architecture among possible alternatives to support DM. This theoretical prediction is consistent with previous research findings based on machine learning techniques(17).

In some cases, e.g. in the delayed response version of the visual motion discrimination task(8), the choice must be maintained actively in working memory. Robust WM requires shielding the interference from both internal noise and external distraction. To assess how the signal-selective structure influences the WM function, a distractor stimulus, whose stimulus strength is equal to the original visual motion stimulus while the contrast condition is inverse(the excitatory population *E*_2_ receives larger external input), is applied to the two excitatory populations. Fig2(a) shows that it takes shorter time to make a different choice(switch to the other decision state) in the presence of the distractor stimulus as the factor *w* is increased. The robustness of the decision attractor can be quantified through the barrier height between the decision attractors that are inferred from landscape topography. As shown in Fig2(b), the barriers separating the current decision attractor from the other one are reduced as the circuit architecture changes from the one with non-selective inhibition to the one with selective inhibition. Our results suggest that the signal-selective structure is less robust against distractors. On the other hand, the weakened stability of the decision state due to the strengthened intermediate state implies that the initial errors are more likely to be corrected during the presence of the visual stimuli. This can further improve the accuracy of DM in addition to the increased accuracy of the initial choices. In summary, there is a tradeoff between the accuracy in DM and the robustness in WM.

**Fig. 2.**
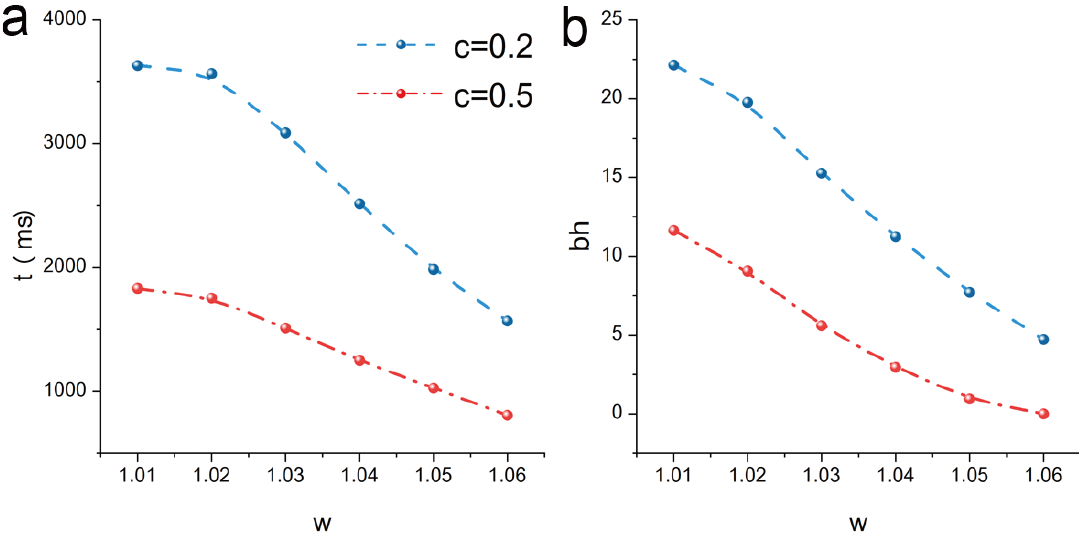
The robustness of WM against distractors. (a)The average transition time from the correct choice to the error one in the presence of distractors for different circuit architectures. The system has to go across a potential barrier to switch to another decision state. The barrier height(bh) inferred from the underlying landscape topography can measure the robustness of the decision state in WM. Both the average transition time and the corresponding barrier heights are reduced as the *w* is decreased, which implies less robustness against distractors.

### Ramping input enhances the stability of working memory

Our results suggest that the circuit architecture with selective inhibition enhances the accuracy in DM while reduces the robustness of the decisions held in working memory to distracting stimuli. This raises an interesting question: what is the underlying mechanism of protecting memory from interference in the circuit with the selective-inhibition architecture? If the circuit architecture is specified, then one probability is that gating incoming stimuli might be achieved by an additional input to the circuit. To test this hypothesis, a non-selective input(the same input to the both selective populations) is applied. We examine how the non-selective input influences the ‘ dynamical behavior and the underlying attractor landscape of the system in DM and WM tasks.

Fig.3(a) shows that the average transition time to the other decision state in the presence of the distracting stimuli increases as the non-selective input is increased. Fig.3(b) shows that the proportion of switching when the transient distracting stimulus is presented in a limited duration(1*s*) reduces with the increased non-selective input. The mechanism of the enhanced robustness against distractors due to the addition non-selective input can be uncovered from the changes in the underlying landscape topography(Fig.3(c)). Both the two decision attractors become stronger with increased non-selective input. The larger non-selective input increases not only the separation between two decision attractors, but also the depth of the attractor basins and the barrier between them(Fig.3(c-f)). It is less likely to go across the barrier and switch to the other decision state in the presence of a larger non-selective input, thereby resulting in the gating of the distracting inputs. These results are consistent with the recent experimental work suggesting that the reduction in sensitivity to distractors results from an external ramping input(35).

**Fig. 3.**
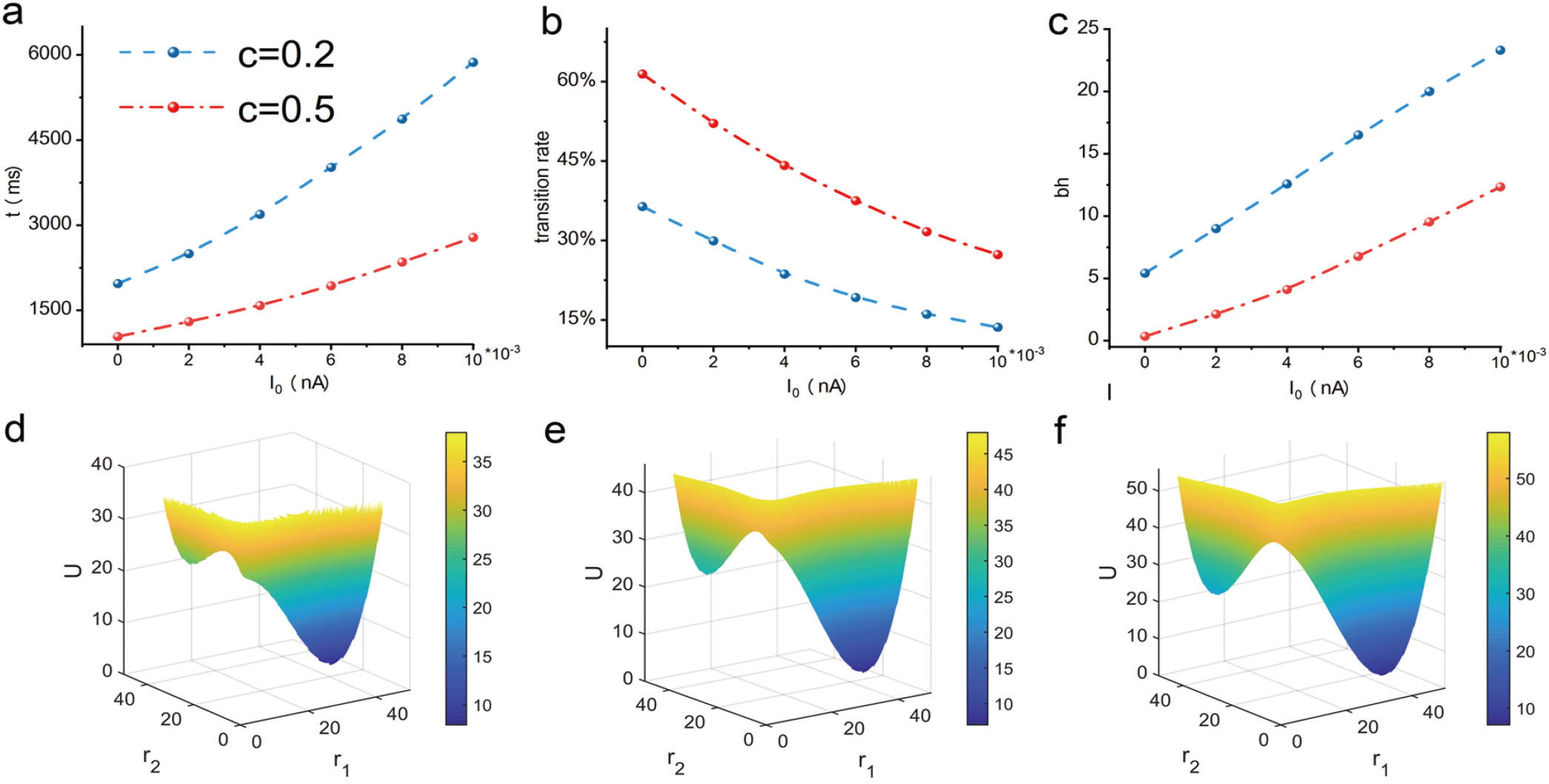
An increasing nonselective input can enhance the robustness of WM against distractors. (a)The average transition time from the correct choice to the error one in the presence of distractors increases with increased nonselective input to both selective excitatory neural populations. (b)The transition rate in a limited duration of the distracting stimulus(1 second) is reduced as the nonselective input is increased. The mechanism of improved robustness is reflected from the larger barrier height(bh) inferred from the underlying landscape topography(Fig.3(c)). The detailed attractor lanscapes with increased nonselective input are displayed in Fig3.(d-f), where the nonselective input *I*_0_ = 0, 0.004, 0.008*nA*, respectively with the specific nonzero contrast *c* = 0.2 and *w* = 1.

Interestingly, the experimental recordings in the previous work(35) show that the distracting stimuli became gradually less capable of affecting the behavior response over time, which implies that a ramping input rather than an input with a constant strength renders the system insensitive to distracting stimuli. Since we have found that larger non-selective input can strengthen the decision states and further result in better stability to distractors, we asked why the brain doesn’t choose to apply a strong non-selective input directly at the beginning of the delay period or even at the beginning of the decision-making tasks? To address this question, we tested the influences of a non-selective input on the DM and WM function from the following three aspects: the speed, accuracy and cost. We first examined the performance in DM tasks with increased non-selective input that is applied from the onset of the visual motion stimuli. Figs.4(a-b) show that the increased non-selective input can lead to faster decisions while worse accuracy. The underlying landscapes that determine these behavioral responses are similar to the ones shown in Fig.3(f). The stronger decision attractors reduce the integration time of the evidence, which results in more errors. This can explain why a strong non-selective input with the constant strength is not applied in the decision process. It can reduce the performance in DM tasks, which seems to be in contradiction with mechanism underlying the circuit architecture with selective inhibition.

**Fig.4.**
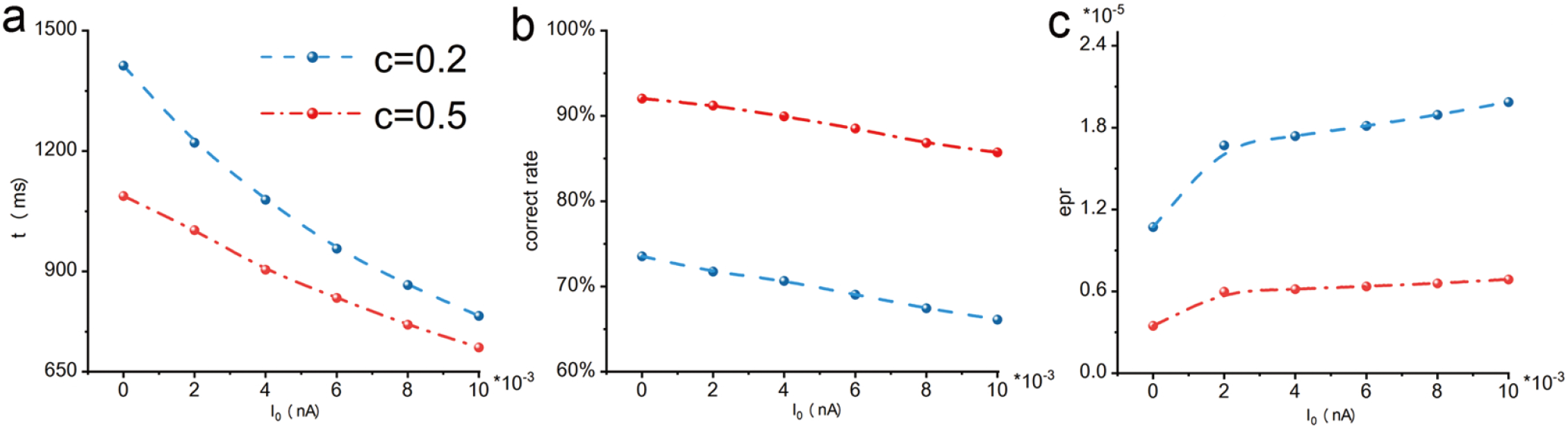
The disadvangates of strong nonselective input in DM and WM.Although the increased nonselective input can improve the robustness of WM against distractors. It leads to shorter decision time(Fig.4(a)) while less accurate choices(Fig.4(b)) and larger energy cost(Fig.4(c)). Presenting A ramping input during the early-delay period may serve as a cost-effective mechanism of temporal gating of distractors

We next explored the mechanism by which an external ramping input is applied in gating distracting stimuli through measuring the thermodynamic cost in maintaining the WM against distractors. Neural circuit as open systems having constant exchange of material,information and energy with the environment, are intrinsically non-equilibrium. In contrast to the steady states in equilibrium systems, the steady states(persistent activities) in neural circuits can only be maintained through constantly consuming energy. The thermodynamics cost for maintaining these activities can be measured by the entropy production rate in non-equilibrium systems (21, 22, 24, 36–38). Fig.4(c) shows that such thermodynamic cost is larger as the applied non-selective input is increased. This result suggests the stability of WM against distractors are enhanced with the expense of more thermodynamic cost. Applying a ramping input serves as the mechanism of gating distractors probably because this is a more cost-effective way. For instance, the system may initially make an incorrect choice, especially for difficult tasks(the contrast *c* is close to 1). Some stimuli presented during the early-delay period may convey valuable information rather than serving as interference signals. These additional stimuli may help to correct the initial errors and improve the accuracy. A ramping input can gradually enhance the robustness of WM, and thus offers an opportunity to improve the performance with as little cost as possible. This mechanism is also consistent with the strategy in forming the circuit architecture that the accuracy is emphasized in DM tasks.

## Discussion

Decision-making (DM) and working-memory(WM) are building blocks of cognition, and great efforts have been made to find out the neural circuit mechanisms of these two functions(3). In the attractor network framework, the characteristic neural activity of WM and DM exhibited in prefrontal cortex and posterior parietal cortex can be reproduced in the circuit architecture with strong recurrent excitation within selective populations and lateral inhibition through a single pool of inhibitory neurons(3, 10, 13). However, recent experimental and modelling evidences suggest that specific functional subnetworks exist within inhibitory populations, which reject the previously proposed circuit mechanisms assuming a single non-selective pool of inhibitory neurons(17). The mechanism underlying DM and WM functions based on such circuit architecture is still unclear.

The attractor landscape can provide a global and quantitative picture describing the dynamics of neural circuit systems. Different landscape topographies indicate that the number and the relative weights of the attractors(functional states) are varied in the underlying landscapes, which determine the categorical choice in decision-making and the robustness of working memories. Although the concept of attractor landscapes has been extensively introduced to describe cognitive functions, they are mostly described as illustrations(3, 10, 13) or only quantified in limited specific circuits(19). Inspired by thermodynamics and statistical mechanics for the physical systems, we developed a non-equilibrium potential landscape and flux framework for general neural circuits. Applying this approach to a biophysically based model that can perform the computations of DM and WM, we explored the circuit mechanisms of DM and WM through focusing on the influences of varied circuit structures on the behavioral responses of the system and the underlying attractor landscapes.

We found that the circuit with the selective-inhibition architecture can support three stable states: one resting state and two decision states in the presence of the visual stimuli.

In contrast to the circuit architecture with non-selective inhibition where only two decision states can be left when the visual stimuli are presented, the stronger resting state result in lengthened time of the integration of evidences. Therefore, the decision time is longer with higher accuracy in the circuit architecture with selective-inhibition. This result is consistent with the prediction in previous results with the machine learning method(17). Protecting the decision held in WM from interference is crucial for cognitive tasks, e.g. a delayed response version of the DM task. However, whether the stability of WM can also be benefited from such circuit architecture is still unknown. Our results suggest that the memories are less robust(shortened transition time to the other decision state) against distractors, which results from the reduced barriers between decision attractors on the underlying landscapes. This result suggests that there is a tradeoff between the DM and WM functions based on a specific circuit architecture. Moreover, this prediction, combined with the experimental observations showing that temporal gating can occur for distractors during DM(35), argues that there may be some additional dynamic modulation mechanisms beyond the circuit architecture for enhancing the performance of DM and WM tasks.

We found that the presence of a ramping non-selective input during the delay period of the DM tasks can serve as a cost-effective mechanism of temporal gating of distractors. A larger nonselective input can induce stronger decision attractors and higher barriers in between. Therefore, it is less probable to change the initial choice by the distractors. However, there is a price for the enhanced robustness of WM. Neural circuits are non-equilibrium systems having constant exchange of materials and energy with the environments. In our non-equilibrium landscape and flux framework, the thermodynamic cost for maintaining the non-equilibrium stable states can be measured in terms of the entropy production rate. Our results show that maintaining the landscapes with stronger decision attractors induced by the additional non-selective input requires more thermodynamic consumption. In other words, better robustness, more thermodynamic cost. Furthermore, the additional non-selective input can lead to shorter decision time while lower accuracy in DM. This would appear to contradict the circuit mechanism that the selective inhibition structure is formed for the purpose of better accuracy in DM. Therefore, the additional non-selective input should only be presented during the delay period against distractors. Moreover, in the early-delay period, some valuable information may be transmitted to the DM circuit. These information are helpful for correcting the errors that could occur in the initial choice and thus should not be gated. Take all of these into account, higher accuracy in DM can be achieved through the circuit architecture with selective inhibition and then a ramping non-selective input presented during the delay period may be the most efficient way to guarantee the correct choice being more robust with less cost.

Sometimes, the working memory system needs to be highly flexible rather than being robust to incoming stimuli according to the environmental conditions or behavioral task demands(15, 39). Previous work has investigated the biophysical mechanisms responsible for the flexibility in terms of the ability to erase memory(back to the resting state) using a negative input(40). A recent study suggests that the intermediate state with high activity for both selective neural populations resulting from the increased recurrent excitation within these neural pools can enhance the flexibility to new stimuli(14). In the present work, we found that the transitions between the different memory states can be promoted through the circuit architecture with selective inhibition where the resting state serves as the intermediate state of the switching. From the perspective of the DM function, such intermediate state increases the accuracy of the initial choices and further may help to correct the initial errors in the presence of the visual stimuli. From the perspective of the WM ability, such intermediate state renders the circuit more flexible to the most recent incoming stimulus. That is, it is easier to load new inputs and discard old memories. This implies a new network mechanism that the enhanced flexibility to new stimuli in WM does not require an addition negative input to reset the system or the the intermediate state resulting from the increased recurrent excitation. The enhanced flexibility can instead arise from the circuit architecture with selective inhibition, which results in strengthened resting state as the intermediate state for transitions to another memory state. Future experiments monitoring the neural activity in subjects trained to remember the latest stimulus will reveal whether the enhanced flexibility originates from the selective inhibition circuit.

Our results imply that the cardinal design principles for circuit architectures supporting DM and WM may be the higher accuracy in DM, accompanied with higher flexibility to incoming signals in WM. Once the robustness against distractors needs to be emphasized according to the behavior task demands, the presence of a ramping non-selective input can reconfigure the dynamics of the circuit and gradually reduce the sensitivity to distractors. This temporal gating mechanism(35) complements the specific functional subnetworks in the circuit architecture for achieving an optimal balance in the tradeoff between the DM and WM functions. Our approach can provide a global and physical quantification which helps to uncover the underlying mechanisms of various biological functions, e.g. a new circuit structure and beyond that for cognitive functions, based on the non-equilibrium physics.

## Materials and Methods

### Model architecture

Here we explored a biophysics-based model that is able to perform WM and DM computations. (3, 10, 41, 42). The model contains two selective, excitatory populations, labeled *E*_1_ and *E*_2_. These two populations have self-excitations from the strong recurrent excitatory connections whith in each excitatory population and the overall effective connectivity between the two excitatory populations is inhibitory through the inhibitory population(3). In contrast to the original models showing that the excitatory populations receive inhibition from a common pool of inhibitory interneurons(3, 10), recent studies suggest subnetworks of selective inhibitory neurons exist in the decision-making circuit(17). We divide the inhibitory neurons into two sub-populations, labeled *I*_1_ and *I*_2_, as shown in Fig.1(b). To manipulate the degree of selectivity of inhibitory neurons, we set the connection strengths between two sub-populations with the same subscript stronger by multiplied a control factor *w,* while the across-population connection strengths smaller by multiplied the factor *w*_ − = 1 – *f* (*w* – 1)*/*(1 – *f*), *f* = 0.45. The factor *w*_ is chosen as the one in the previous work describing how the synaptic weights between neurons belonging to different clusters were depressed (43). The value of *w* determines how selective the inhibitory neurons are: *w* = 1 corresponds to the nonselective inhibition while larger *w* implies more selective inhibition.

Previous modelling works suggest the decision-making circuit that consists of thousands spiking neurons can be reduced into a two-variable model through several approximations, e.g. linearization of the input–output relation of the inhibitory interneurons(3, 10). Based on the fact that the synaptic gating variable of NMDA receptors has a much longer decay time constant than other timescales in the circuit, the dynamics of the system can be described by the dynamical equations of the average NMDA synaptic gating variable 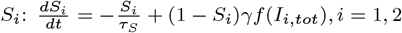, where firing rate *r_i_* of neural population *i* is a function of total input current *I_i,tot_* that can be written as(10, 42): 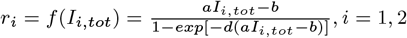. The corresponding parameters are *a* = 270(*V_n_C*)^-1^, *b* = 108Hz, *d* = 0.154*s*, *γ* = 0.641 and *τ_S_* = 100*ms*, the same with the ones shown in the previous work(10). The total synaptic input currents of the two neural populations dominated by the NMDA receptors are:

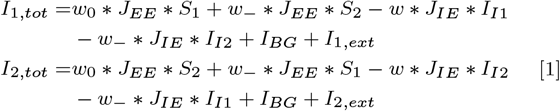

here *J_EE_* is the effective coupling constant between excitatory subpopulation and *J_IE_* is the coupling constant from the inhibitory sub-population to the excitatory sub-population for the non-selective case(*w* = 1). The factor *w*_0_ = 1.7 indicates the neurons belonging to the same excitatory sub-population with potentiated synaptic weight. With the linear approximation of the input–output transfer function of the inhibitory cell, the inhibitory input from an inhibitory sub-population to an excitatory sub-population depends on the total synaptic input current(*I*_*I*1_ or *I*_*I*2_) that this inhibitory sub-population receives, where

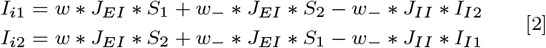

Substitute the Eq. (2) into Eq. (1), we can obtain the total synaptic input currents *I*_1,*tot*_, *I*_2,*tot*_ that only depend on the average NMDA synaptic gating variable *S*_1_ and *S*_2_. It can be easily proven that the corresponding parameters in the input–output transfer function of the inhibitory cell can be absorbed into the coupling constants *J_IE_, J_EI_, J_II_* and the background input *I_BG_*. Here the particular values of the parameters are set as *J_EE_* = 0.48*nA, J_II_* = 5*nA, J_EI_* = 0.52*nA, J_IE_* = 1*nA, I_BG_* = 0.31*nA* based on the references(3, 10, 17).

To characterize the DM function in this circuit model, the external stimulus input to the selective neural population is introduced as: *I_i,ext_* = *J_A,ext_* * *μ* * (1 ± *c*), *i* = 1, 2, where *J_A,ext_* = 5.2 * 10^-4^*nA* · *Hz*^-1^, *μ*_0_ = 30*Hz* is the external average synaptic coupling with the AMPA receptors. The + or sign refers to whether the stimulus is the preferred one or non-preferred of the corresponding selective neural population. As introduced in the main text, The contrast *c* which sets the bias of the input for one population over the other. For a zero-contrast stimulus(*c* = 0), the two excitatory populations receive equal input. The WM function is characterized by the stimulus-selective persistent activities and their robustness against distractors. The distractor-related current is stimulated as the input with the equal strength to the original external stimulus while the contrast condition is inverse(the excitatory population that initially receives larger external input now receives the smaller input).

The dynamical trajectories of the system are calculated using the Runge-Kutta method with an integration time step of 0.02*ms*. In each trial, the random fluctuation is introduced by a noise term implemented as an uncorrelated standard Gaussian noise with zero mean and the variance equals to 0.025*nA*. For the computations of the average decision time/transition time and correct rate, each data point is obtained from more than 30000 trials.

### Non-equilibrium landscape and flux framework for general neural circuits

The attractor landscape metaphor is widely used to describe cognitive functions such as associative memory retrieval, classification, et al(1, 19, 44). However, such attractor landscape can only be explicitly quantified in limited specific networks, e.g. the symmetrical neural circuit in the original Hopfield model(1, 19). To address the issues of global stability, we developed a non-equilibrium potential landscape and flux theory for the general neural networks(23). We focused on the probabilistic evolution of the neural network rather than following the individual trajectories for characterizing the dynamics globally. Due to the stochastic nature of a realistic neural network, we can describe the stochastic dynamics of the neural network with a set of Langevin equations: 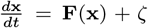. *ζ* represents the stochastic fluctuations, which are assumed to follow a Gaussian distribution with autocorrelations specified by < *ζ*(**x**, *t*)*ζ*(**x**, *t′*) >= 2**D**(**x**)*δ*(*t* – *t′*). Here **D**(**x**) is the diffusion coefficient matrix giving the magnitude of the fluctuations. *δ*(*t*) is a delta function. These stochastic differential equations can be mapped onto an equivalent Fokker-Planck equation for the probability density: *∂P*(**x**, *t*)/*∂t* = –▽· (**F**(**x**)* *P*(**x**, *t*)) + ▽ · (▽ · (**D***P*(**x**, *t*))). Furthermore,the steady state probability distribution *P_ss_* can be solved from ▽ · (**F**(**x**) * *P_ss_*(**x**)) – ▽ · (▽ · (**D***P_ss_*(**x**))) = 0. And then we can quantify the potential landscape by the relationship *U* = –*InP_ss_* (**x**) analogous to equilibrium statistical mechanics where the potential landscape is related to the equilibrium distribution through the Boltzman law.

A distinguishing feature of non-equilibrium systems is that the corresponding driving force is determined by not only a gradient of the underlying potential landscape but also a nonvanishing steady-state flux:**F** = **J***_ss_*/*P_ss_* – **D** · ▽*U* (23, 24, 34, 45, 46). Such non-equilibrium flux signifies the violation of detailed balance. Biological systems, such as neural circuits, consume energy to perform different vital functions(34, 47). The flux as an indispensable part of the non-equilibrium driving force contributes directly to the thermodynamic cost for maintaining the function of the neural network in terms of the entropy production rate. The energy cost computed in this way has been successfully used to explore cost-performance trade-off in biological systems whose energy sources are ATP, GTP and SAM(21, 22, 36).

## ACKNOWLEDGMENTS

We thank Kun Zhang and Li Xu for valuable discussions and suggestions related to this manuscript. H.Y. is grateful for support via National Natural Science Foundation of China Grants 21721003.

## References

1. JJ Hopfield, Neural networks and physical systems with emergent collective computational abilities. Proc. Natl.Acad. Sci. United States Am. Sci. 79, 2554–2558 (1982).

2. TS Deisboeck, JY Kresh, Complex Systems Science in Biomedicine. (Springer US), (2006).

3. JD Murray, J Jaramillo, XJ Wang, Working memory and decision-making in a frontoparietal circuit model. J. Neurosci. 37, 12167–12186 (2017).

4. JP Fadok, et al., A competitive inhibitory circuit for selection of active and passive fear responses. Nature 542, 96–+ (2017).

5. Y Yoshimura, EM Callaway, Fine-scale specificity of cortical networks depends on inhibitory cell type and connectivity. Nat. Neurosci. 8, 1552–1559 (2005).

6. ZQ Lin, CC Nie, YF Zhang, Y Chen, TM Yang, Evidence accumulation for value computation in the prefrontal cortex during decision making. Proc. Natl. Acad. Sci. United States Am. 117, 30728–30737 (2020).

7. JD Schall, Neural basis of deciding, choosing and acting. Nat. Rev. Neurosci. 2, 33–42 (2001).

8. MN Shadlen, WT Newsome, Neural basis of a perceptual decision in the parietal cortex (area lip) of the rhesus monkey. J. Neurophysiol. 86, 1916–1936 (2001).

9. JI Gold, MN Shadlen, The neural basis of decision making. Annu. Rev. Neurosci. 30, 535–574 (2007).

10. KF Wong, XJ Wang, A recurrent network mechanism of time integration in perceptual decisions. J. Neurosci. 26, 1314–1328 (2006).

11. DJ Amit, N Brunel, Model of global spontaneous activity and local structured activity during delay periods in the cerebral cortex. Cereb. Cortex 7, 237–252 (1997).

12. XJ Wang, Synaptic reverberation underlying mnemonic persistent activity. Trends Neurosci. 24, 455–463 (2001).

13. XJ Wang, Attractor network models. Encycl. Neurosci. pp. 667–679 (2009).

14. H Yan, J Wang, Non-equilibrium landscape and flux reveal the stability-flexibility-energy tradeoff in working memory. PLoS Comput. Biol. 16, e1008209 (2020).

15. CK Machens, R Romo, CD Brody, Flexible control of mutual inhibition: A neural model of two-interval discrimination. Science 307, 1121–1124 (2005).

16. XJ Wang, Probabilistic decision making by slow reverberation in cortical circuits. Neuron 36, 955–968 (2002).

17. F Najafi, et al., Excitatory and inhibitory subnetworks are equally selective during decision-making and emerge simultaneously during learning. Neuron 105, 165–+ (2020).

18. JD Roitman, MN Shadlen, Response of neurons in the lateral intraparietal area during a combined visual discrimination reaction time task. J. Neurosci. 22, 9475–9489 (2002).

19. JJ Hopfield, DW Tank, Computing with neural circuits - a model. Science 233, 625–633 (1986).

20. PG Wolynes, JN Onuchic, D Thirumalai, Navigating the folding routes. Science 267, 1619–1620 (1995).

21. Y Cao, H Wang, Q Ouyang, Y Tu, The free-energy cost of accurate biochemical oscillations. Nat. physics 11, 772 (2015).

22. G Lan, P Sartori, S Neumann, V Sourjik, Y Tu, The energy “cspeed”caccuracy trade-off in sensory adaptation. Nat. physics 8, 422 (2012).

23. H Yan, et al., Nonequilibrium landscape theory of neural networks. Proc. Natl. Acad. Sci. United States Am. 110, E4185–E4194 (2013).

24. J Wang, Landscape and flux theory of non-equilibrium dynamical systems with application to biology. Adv. Phys. 64, 1–137 (2015).

25. J Duncan, The multiple-demand (md) system of the primate brain: mental programs for intelligent behaviour. Trends Cogn. Sci. 14, 172–179 (2010).

26. DJ Mitchell, et al., A putative multiple-demand system in the macaque brain. J. Neurosci. 36, 8574–8585 (2016).

27. P Domenech, J Redoute, E Koechlin, JC Dreher, The neuro-computational architecture of value-based selection in the human brain. Cereb. Cortex 28, 585–601 (2018).

28. WT Newsome, KH Britten, JA Movshon, Neuronal correlates of a perceptual decision. Nature 341, 52–54 (1989).

29. MN Shadlen, WT Newsome, Motion perception: Seeing and deciding. Proc. Natl. Acad. Sci. United States Am. 93, 628–633 (1996).

30. A Resulaj, R Kiani, DM Wolpert, MN Shadlen, Changes of mind in decision-making. Nature 461, 263–U141 (2009).

31. GM Stine, A Zylberberg, J Ditterich, MN Shadlen, Differentiating between integration and non-integration strategies in perceptual decision making. Elife 9 (2020).

32. N Brunel, XJ Wang, Effects of neuromodulation in a cortical network model of object working memory dominated by recurrent inhibition. J. Comput. Neurosci. 11, 63–85 (2001).

33. CC Lo, CT Wang, XJ Wang, Speed-accuracy tradeoff by a control signal with balanced excitation and inhibition. J. Neurophysiol. 114, 650–661 (2015).

34. H Yan, K Zhang, J Wang, Physical mechanism of mind changes and tradeoffs among speed, accuracy, and energy cost in brain decision making: Landscape, flux, and path perspectives. Chin. Phys. B 25 (2016).

35. A Finkelstein, et al., Attractor dynamics gate cortical information flow during decision-making. Nat. Neurosci. 24, 843–+ (2021).

36. L Xu, H Shi, H Feng, J Wang, The energy pump and the origin of the non-equilibrium flux of the dynamical systems and the networks. The J. chemical physics 136, 04B621 (2012).

37. H Ge, H Qian, Dissipation, generalized free energy, and a self-consistent nonequilibrium thermodynamics of chemically driven open subsystems. Phys. Rev. E 87, 062125 (2013).

38. H Yan, B Li, J Wang, Non-equilibrium landscape and flux reveal how the central amygdala circuit gates passive and active defensive responses. J. Royal Soc. Interface 16 (2019).

39. JD Cohen, et al., Temporal dynamics of brain activation during a working memory task. Nature 386, 604–608 (1997).

40. J Pereira, XJ Wang, A tradeoff between accuracy and flexibility in a working memory circuit endowed with slow feedback mechanisms. Cereb. Cortex 25, 3586–3601 (2015).

41. HR Wilson, JD Cowan, Excitatory and inhibitory interactions in localized populations of model neurons. Biophys. journal 12, 1–24 (1972).

42. G Deco, et al., Resting-state functional connectivity emerges from structurally and dynamically shaped slow linear fluctuations. J. Neurosci. 33, 11239–11252 (2013).

43. L Mazzucato, G La Camera, A Fontanini, Expectation-induced modulation of metastable activity underlies faster coding of sensory stimuli. Nat. Neurosci. 22, 787–+ (2019).

44. DJ Amit, Modeling brain function: The world of attractor neural networks. (Cambridge University Press), (1992).

45. J Wang, L Xu, EK Wang, Potential landscape and flux framework of nonequilibrium networks: Robustness, dissipation, and coherence of biochemical oscillations. Proc. Natl. Acad. Sci. United States Am. 105, 12271–12276 (2008).

46. J Wang, K Zhang, L Xu, E Wang, Quantifying the waddington landscape and biological paths for development and differentiation. Proc. Natl. Acad. Sci. United States Am. 108, 8257–8262 (2011).

47. E Eisenberg, TL Hill, Muscle contraction and free energy transduction in biological systems. Science 227, 999–1006 (1985).

